# Interference with HIV infection of the first cell is essential for viral clearance in a pre-exposure prophylaxis model

**DOI:** 10.1101/435552

**Authors:** Ana Moyano De Las Muelas, Gila Lustig, Hylton E. Rodel, Tibor Antal, Alex Sigal

**Author notes:** For correspondence (AS).

## Abstract

Pre-exposure prophylaxis (PrEP) uses relatively weak HIV inhibition to reduce transmission between individuals. Why this approach is successful is unclear. Here we derive and experimentally validate a mathematical model for predicting infection clearance with PrEP based on the measured effect of a drug on the HIV replication ratio and number of initial infected cells. We tested the model by inhibiting low dose HIV infection with tenofovir, which reduces infection frequency per cell, and atazanavir, which reduces the cellular burst size of viable virions. Both drugs were at concentrations which allowed similar HIV replication. Reducing infection frequency dramatically increased infection clearance, while reducing burst size did not. This indicates that initial infection is vulnerable to inhibition before it infects the first cell, but not thereafter. Our model explains why PrEP is potent at drug concentrations which are ineffective against established infection, and provides a framework to test drug effectiveness for PrEP.

## Introduction

Antiretroviral therapy (ART) currently used to treat ongoing HIV infection combines several potent antiretroviral drugs (ARVs). Yet, no current ART regimen has been successful in extinguishing infection once it is established in the human host (*Churchill et al.* (*2016*); *Sigal and Baltimore* (*2012*)). In contrast to the robustness of HIV infection once established, initial HIV transmission between an infected and uninfected individual is more fragile. The probability for an exposed individual to become infected does not exceed 0.02 per sexual act under any set of conditions and is usually much lower (*Gray et al.* (*2001*); *Jin et al.* (*2010*)). It generally involves one founder viral clone (*Keele et al.* (*2008*)). This indicates that an infection bottleneck exists, where most infection attempts by transmitted virions are unsuccessful.

The probability of initial transmission has been shown to be effectively reduced with a single ARV if the ARV is present before exposure takes place, an approach termed PrEP. The ARVs used for PrEP are a subset found in the full treatment regimen. A central agent is tenofovir (TFV), which interferes with viral reverse transcription and is usually delivered as the prodrug tenofovir disoproxil fumarate (TDF) or tenofovir alafenamide fumarate (TAF TFV is either used by itself or co-administered with emtricitabine (FTC), another HIV reverse transcriptase inhibitor. This approach was shown to be effective in a non-human primate model of low dose infection, even when dosing was intermittent (*García-Lerma et al.* (*2008*)). In contrast to PrEP, TFV and FTC are only used in combination with a highly potent ARV such as dolutegravir or efavirenz to suppress established infection (https://aidsinfo.nih.gov/guidelines/html/1/adult-and-adolescent-arv/11/what-to-start).

**Figure 1.**
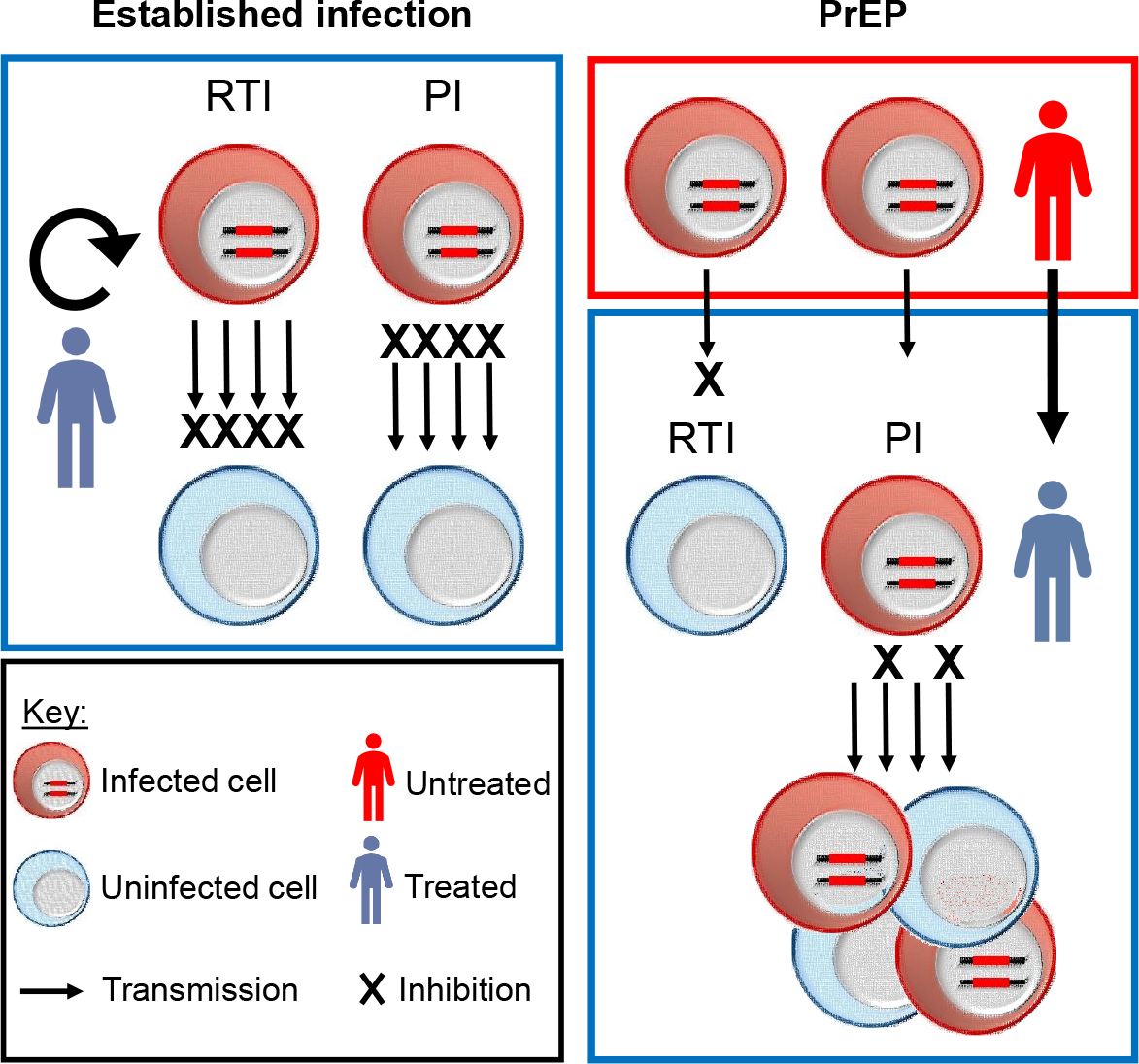
Inhibition of established versus transmitted infection. Left panel shows established infection in an individual, where both the infected donor cell and uninfected target cell are exposed to antiretroviral drugs. In this case, a complete block of transmission is illustrated, with infection persisting in a reservoir of long lived infected cells. Right panel shows infection in PrEP, where the virus is produced in an infected individual where suppression of infection by drugs is absent or ineffective. The virus is then transmitted to an uninfected, PrEP treated individual. Two drug mechanisms of inhibition are shown: Reverse transcriptase inhibitors (RTI) and other ARVs such as integrase inhibitors reduce infection frequency by blocking HIV from becoming a provirus in the host DNA capable of making new infectious virions. Protease inhibitors (PI) reduce the burst size of infectious virions per infected cell by interfering with HIV maturation, a process which requires cleavage of HIV proteins by the HIV protease.

The majority of clinical studies have shown PrEP effectiveness in a variety of populations, transmission modes, and drug delivery modalities (*Baeten et al.* (*2012*); *Choopanya et al.* (*2013*); *McCormack et al.* (*2016*); *Molina et al.* (*2015*); *Thigpen et al.* (*2012*); *Grant et al.* (*2010*); *Karim et al.* (*2010*); *Baeten et al.* (*2016*); *Nel et al.* (*2016*)). These include studies utilizing oral TDF alone (*Baeten et al.* (*2012*); *Choopanya et al.* (*2013*)), oral TDF/FTC (*Baeten et al.* (*2012*); *McCormack et al.* (*2016*); *Molina et al.* (*2015*); *Thigpen et al.* (*2012*); *Grant et al.* (*2010*)), TFV gel (*Karim et al.* (*2010*)), and vaginal rings containing the reverse transcriptase inhibitor dapivirine (*Baeten et al.* (*2016*); *Nel et al.* (*2016*)). Currently, PrEP is indicated when there is a high likelihood of HIV infection due to sexual contact with an HIV infected individual or injectable drug use (*WHO* (*2015*); *Anderson et al.* (*2012*)) and may be used in conjunction with other prevention measures to reduce HIV incidence (*Punyacharoensin et al.* (*2016*); *Pretorius et al.* (*2010*)). The reason to use a weakened ARV regimen is partly to decrease side effects and thus make it more acceptable to healthy individuals (*Choopanya et al.* (*2013*); *Koechlin et al.* (*2017*)).

Why weak inhibition relative to the replicative capacity of HIV is effective in PrEP has not been investigated theoretically or experimentally, and such understanding may guide the development of new PrEP approaches. Currently approved ARVs include drugs which target various stages of the viral life cycle (http://aidsinfo.nih.gov/). Among the most commonly used are inhibitors of the HIV proteins reverse transcriptase, integrase, and protease. Reverse transcriptase and integrase inhibitors prevent the initial infection of the cell but do not interfere with viral production from an already infected cell. Protease inhibitors do not interfere with cellular infection but reduce the number of viable virions an infected cell produces by interfering with virion maturation, which requires the HIV protease. Hence, the two mechanisms of inhibition involve either decreasing HIV infection frequency or viral burst size (*Delbrück* (*1945*)) per infected cell. However, it has been reported that protease inhibitors decrease infection frequency to some extent (*Rabi et al.* (*2013*)).

The effect of decreasing infection frequency or burst size should be symmetrical. The number of successful infections will be decreased if virions are prevented from infecting a cell by blocking their ability to become a provirus capable of generating new virions. It will also be decreased by reducing the output of new viable virions from an infected cell. In either case, fewer virions are available for the next cycle of infection (Figure 1, left panel). Whether or not infection expands is determined by the replication ratio *R*_0_, which measures the mean number of new cells infected by a single infected cell. Given that *R*_0_ = *rκ*, where *r* is the probability of infection and *κ* the viral burst size per cell (*Ribeiro et al.* (*2010*); *Nowak and May* (*2000*)), decreasing *R*_0_ should be possible by either decreasing *r* or *κ*. The exception to the symmetrical effects of the two drug mechanisms on HIV replication is when the virus originates at low numbers in a different individual. In this case, PrEP is applied to the yet uninfected, exposed individual. In HIV infection, if transmission is by cell-free virus (*Jackson et al.* (*2018*); *Boullé et al.* (*2016*); *Sigal et al.* (*2011*); *Kim et al.* (*2018*)), using a drug which reduces burst size may not affect the incoming virus in the first infection cycle, since that virus is already mature (*Arts and Hazuda* (*2012*)). In contrast, a regimen which decreases infection frequency may be effective in preventing the incoming virus from infecting the first cell in the new host (Figure 1, right panel). After initial infection, both inhibitors should act symmetrically. Whether this difference between inhibitor mechanisms is important in determining the clearance probability – the probability that infection will be terminated as a consequence of the inhibitor – is unknown.

Here we modeled HIV infection as a function of the initial mean number of infected cells *λ* and the replication ratio *R*_0_, both of which we measured experimentally. We used two types of inhibition: reduction of infection frequency mediated by TFV, and reduction of the burst size of viable virions mediated by atazanavir (ATV), a drug belonging to the class of HIV protease inhibitors. We observed that while both drugs reduced *R*_0_ to a similar extent at the concentrations used, only TFV, which prevented successful infection of the first set of cells, was effective at clearing infection.

## Results

### Derivation of a model for infection clearance

We first set out to model the effect of reducing infection frequency and burst size on the probability to clear infection (*P*_*c*_). We model the infection chain of HIV transmissions between cells by using a branching process (*Athreya et al.* (*2004*)). We consider the initial stages of HIV infection, where host cells are not limiting in the number of cells that can be infected (*Ribeiro et al.* (*2010*); *Nowak and May* (*2000*)). An infected cell produces a burst of *κ* virions. *κ* is on the order of 10^3^ to 10^4^ (*Eckstein et al.* (*2001*); *Chen et al.* (*2007*)). The probability of one virion infecting the next cell is *r*, the infection frequency. We note the maximum number of cells which can be infected per burst is *κ*. A key quantity is the product *R*_0_ = *rκ* (*Nowak and May* (*2000*)). *R*_0_ is approximately 10 *in vivo* (*Ribeiro et al.* (*2010*)). Eventual infection clearance is certain for *R*_0_ ≤ 1. For *R*_0_ > 1, infection may still be cleared if, at any point in the infection chain, the number of infected cells is zero. Hence, we define the probability that infection is cleared as:

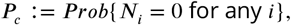

where *N*_*i*_ is the number of infected cells in the *i*-th transmission step.

We denote by *q* the probability that infection starting from a single infected cell is cleared. In the biologically relevant case where *κ* is large and *r* is small, with *R*_0_ finite, we can calculate *q* as (see derivation in Materials and Methods):

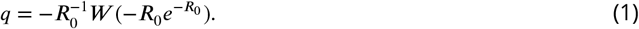

Here, *W* is the Lambert *W*–function (*Gradshteyn and Ryzhik* (*2014*)), the inverse of the function *g*(*x*) = *xe*^*x*^.

As defined, *q* is the probability that the infection originating in a single infected cell will be cleared. We note that in the case where host cells are not limiting, as occurs in the initial stages of infection, infection chains originating from individual infected cells are independent of each other. Hence, given *n* initial infected cells,

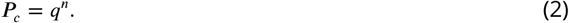

We consider the initial number of infected cells as a random variable from a distribution *ϕ*_*n*_, where *p*_*c*_ = ∑_*n*≥0_ *ϕ*_*n*_ *q*^*n*^. A biologically relevant choice in viral infection is the Poisson distribution *ϕ*_*n*_ = *λ*^*n*^ *e*^−*λ*^/ *n*!, where *λ* is the mean number of initial infected cells. Using *e*^*x*^ = ∑_*n*≥0_ *x*^*n*^/ *n*!, ∑_*n*≥0_ *ϕ*_*n*_ *q*^*n*^ then becomes:

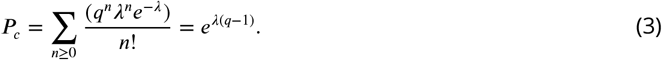

We now consider the effect of the antiretroviral drug mechanism on *λ* and *q*. We note that ARVs reduce either infection frequency *r* or burst size *κ*. For drugs which reduce infection frequency, *r* → *d*_1_*r*, and for drugs which reduce viral burst size, *κ* → *d*_2_*κ*, where 0 ≤ *d*_1_, *d*_2_ ≤ 1. Given *R*_0_ = *rκ* and therefore *R*_0_ → *R*_0_*d*_1_*d*_2_, the effects of the drug mechanisms are symmetrical on *q*:

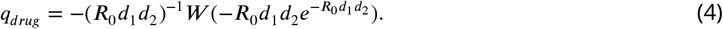

Hence, if the drugs decrease *R*_0_ to a similar extent, their effect on *q* will also be similar. However, given an initial transmission with cell-free virus, only the drug mechanism that decreases infection frequency will reduce the mean initial number of infected cells *λ*. The mechanism which reduces burst size will only affect the success of the next transmission cycle. Therefore, the probability to clear infection with ARVs becomes:

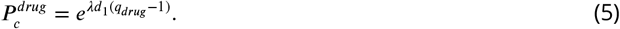

Here *q*_*drug*_ is determined by Eq. (4). Given that *λ* is expected to be small, on the order of one infected cell, the probability of clearance is expected to be very sensitive to the drug mechanism which reduces infection frequency (*d*_1_), provided the effect of the drug is present when initial infection takes place. The drug mechanism which reduces burst size (*d*_2_) misses this initial intervention point.

To visualize the effects of decreasing *R*_0_ versus *λ*, we plotted Eq. (3) for a range of parameter values (Figure 2A). It can be observed that for *R*_0_ ≤ 1, infection terminates. At *R*_0_ ⪆ 1.5, infection is not strongly sensitive to the exact *R*_0_ value. However at all *R*_0_ > 1 values, the probability of infection clearance is very sensitive to *λ*, provided *λ* is small. This sensitivity is greatly reduced when *λ* ⪆ 2.

To examine the effects of drug mechanism, we plotted infection clearance according to Eq. (5) at two conditions of *R*_0_ and *λ* relative to *d*_1_ and *d*_2_ (Figure 2B). In the first condition, *R*_0_ was suZciently small to be decreased below 1 by the drugs in the inhibition range used, while *λ* was large. In the second condition, *R*_0_ was large while *λ* small. In the first condition, both drug mechanisms had a similar effect on infection clearance, and *P*_*c*_ = 1 when the effect of either drug reduced *R*_0_ below 1. In the second condition, only *d*_1_, which decreased infection frequency, substantially increased *P*_*c*_. *d*_2_, which acted on burst size, had no appreciable effect. We note that based on observations of *R*_0_ ≈ 10 *in vivo* (*Ribeiro et al.* (*2010*)) and a probability of clearance of at least 0.98 per exposure from an HIV positive partner in the absence of PrEP (*Gray et al.* (*2001*); *Jin et al.* (*2010*)) which is suggestive of a small value for *λ*, the second condition likely reflects the physiological situation.

**Figure 2.**
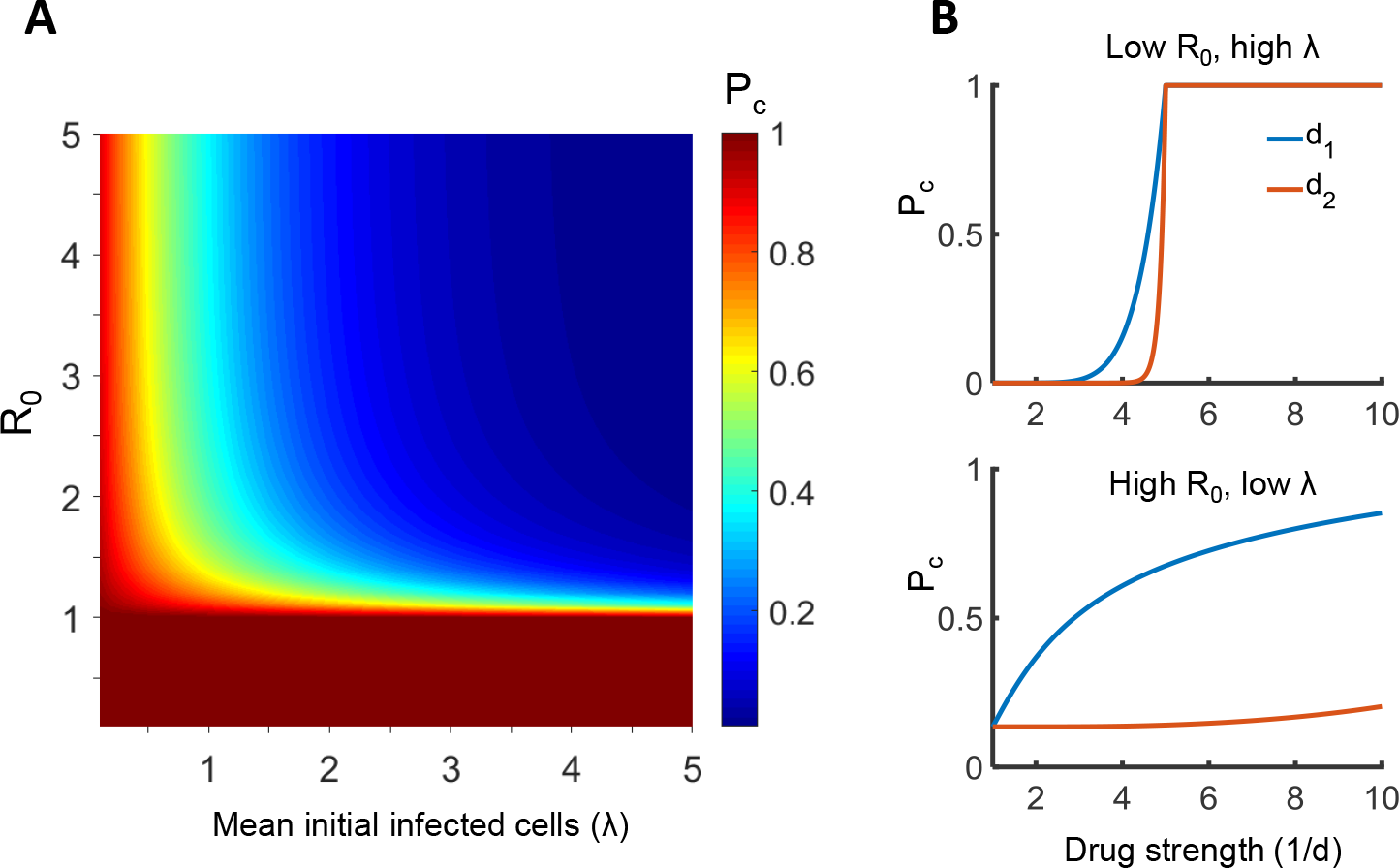
Effects of the initial infected cell number *λ* and replication ratio *R*_0_ on the probability of infection clearance *P*_*c*_. (A) *P*_*c*_ according to Eq. (3) at different parameter values for *λ* and *R*_0_. (B) *P*_*c*_ according to Eq. (5) when a drug attenuating infection frequency (*d*_1_) or burst size (*d*_2_) acts on *λ* and *R*_0_. Top panel shows the case where *R*_0_=5, *λ* =20, while bottom panel shows the case where *R*_0_.=20, *λ* =2. X-axis is drug strength as 1/*d*, y-axis is *P*_*c*_

### Experimental determination of the probability of infection clearance

We examined experimentally whether Eq. (5) predicts *P*_*c*_ for different drug mechanisms at a low number of initial infected cells, the likely *in vivo* condition for initial infection. We used the reverse transcriptase inhibitor TFV and the protease inhibitor ATV to inhibit infection initiating as cell-free HIV. We measured the effect of each drug on the initial number of infected cells *λ* from cell-free infection and on *R*_0_ after the first infection cycle. For virus, we used HIV NL(AD8), an HIV strain with a CCR5 tropic envelope protein. CCR5 tropism has been shown to be the predominant transmitted form between individuals (*Keele et al.* (*2008*)). As target cells for infection, we used a clone of the RevCEM infection indicator cell line (*Wu et al.* (*2007*)) which we first subcloned to increase detection efficiency (*Boullé et al.* (*2016*)) then modified to express the CCR5 receptor (RevCEM-B8, Materials and Methods). Detection of infected cells was done by detecting the number of GFP positive cells using flow cytometry.

We titrated TFV and ATV to obtain a similar effect on ongoing viral replication, which occurred at 60 *μ*M and 16 *n*M of TFV and ATV respectively. We then infected cells in the presence of these drug concentrations and measured infection over time to precisely measure *R*_0_ for each drug condition. We initiated infection with the same dose of cell-free HIV for each infection condition. After initiation of infection, ongoing infection was maintained by transmission through coculture of infected and uninfected cells. To maintain nutrients for cell growth and prevent the depletion of the uninfected target cell population, we passaged cells every two days by diluting infection with fresh medium at a 1:2 ratio for infection conditions with drug. Proliferation of uninfected cells was suZcient to maintain uninfected cell numbers, and infection was below 5 percent for both drug conditions at all time-points, ensuring target cells were not limiting (Figure 3A). For the no drug condition, the infection expanded much more rapidly. Therefore, the infected cell culture was passaged by diluting the infected cells 1:100 every 2 days into uninfected cells. Infection was monitored over 8 days. Despite the use of the same HIV cell-free input dose, there were pronounced differences at day 2 between TFV and ATV (Figure 3A). This time-point reflects the results of the initial cell-free infection given an approximately 2 day viral cycle (*Perelson et al.* (*1996*)). Cell-free infection was strongly inhibited by TFV relative to no drug. In contrast, the effect of ATV on cell-free infection was much weaker. After the day 2 time-point, infection expanded with similar dynamics for both drug conditions, and much more rapidly when no drug was present.

**Figure 3.**
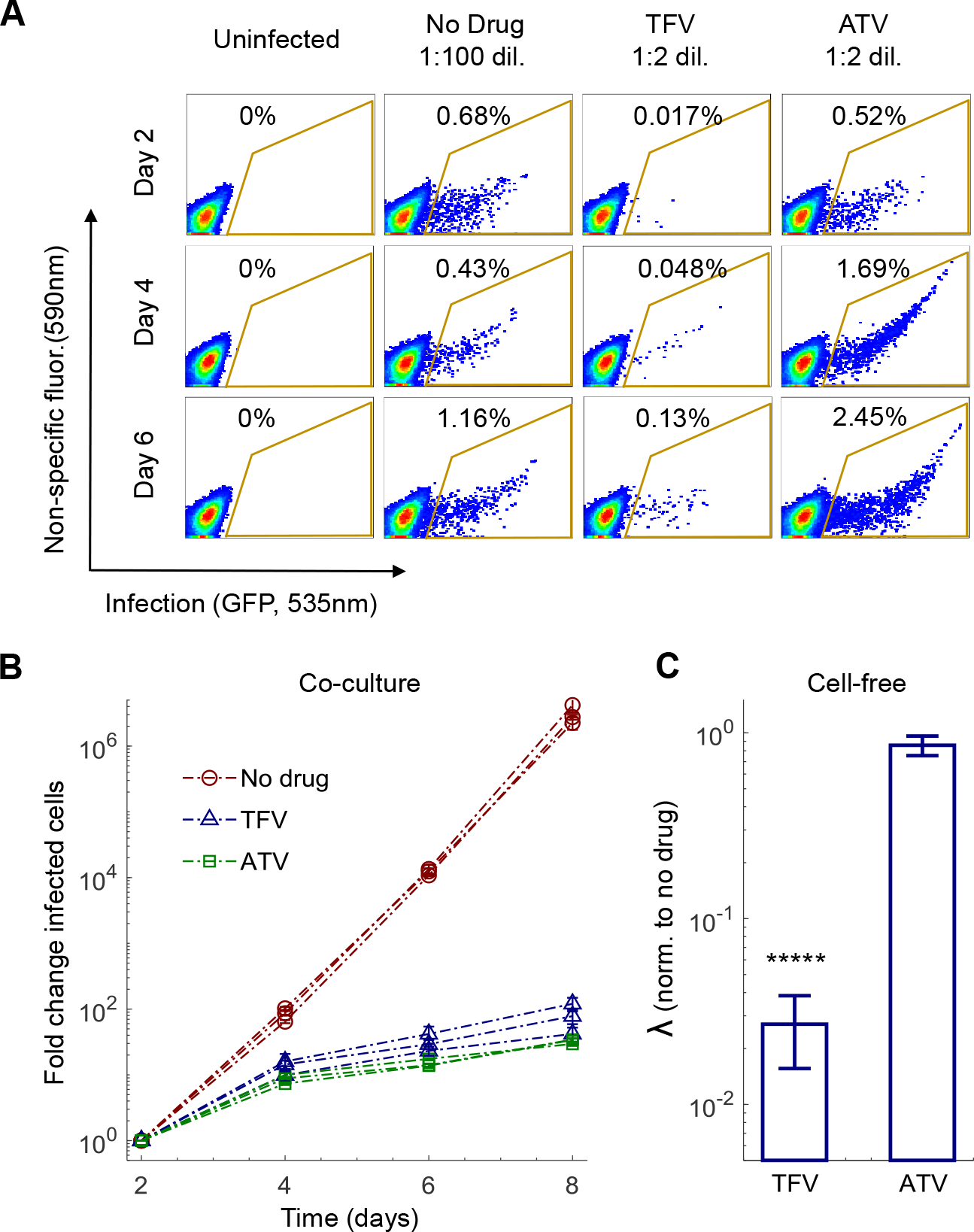
Experimental measurement of drug effect on *R*_0_ and the initial number of infected cells *λ*. (A) Flow cytometry plots of the fraction of infected cells at different days post infection in the absence of drug or presence of 60 *μ*M TFV or 16 *n*M ATV. Day 2 is the first time-point after the initial cell-free infection, corresponding to approximately one viral cycle. X-axis is GFP fluorescence, y-axis is autofluorescence, with the fraction of infected cells corresponding to the cells within the area outlined in yellow. Infected cell cultures in the presence of either drug were diluted 1:2 every 2 days. Infected cultures in the absence of drug were diluted 1:100 into uninfected cells every 2 days. (B) Measurement of *R*_0_ in the absence and presence of drug. The number of infected cells at each time-point is normalized by the number of infected cells at day 2 and corrected for the dilution factor used in each infection cycle. 3 independent experiment were performed, with each point denoting the mean ± std of 3 experimental replicates per experiment. Infection in the absence of drug is shown as red circles, TFV as blue triangles, and ATV as green squares. (C) Effect of drug on *λ*. For each drug condition *λ* was measured 2 days after cell-free HIV infection and normalized by *λ* for no drug. Mean ± std of 3 independent experiments, where normalization was with *λ* in the absence of drug as measured in the same experiment. Raw numbers of infected cells averaged over all experiments were 1.3*x*10^4^ ± 1.5*x*10^3^ for no drug infection, 3.4*x*10^2^ ± 1.3*x*10^2^ for TFV and 1.1*×*10^4^ ± 7.4*x*10^3^ for ATV (mean ± std).

We plotted the total number of infected cells, corrected for cells removed during passaging, versus time (Figure 3B). We then calculated the effect of drug on *R*_0_ over a two day cycle (Table 1). *R*_0_ values were calculated as 4.2 ± 0.73 for TFV and 3.2 ± 0.088 for ATV (mean ± std). *R*_0_ for infection in the absence was 143 ± 15 in the cell line. We then measured the effect of the drugs on *λ* after the first cycle of infection (day 0 to day 2), and compared the results to infection in the absence of drug. *λ* in the presence of drug divided by *λ* for the no drug condition (*λ*_*norm*_) was 0.027 ± 0.014 for TFV and 0.88 ± 0.16 for ATV, (Figure 3C, Table 1). The decrease in *λ* for TFV versus ATV was significant (p=6*x*10^−14^, t-test).

**Table 1.** Measured Parameter Values

| Treatment | $\lambda_{norm}$ | $R_0$ |
| --- | --- | --- |
| No drug | 1 | $143 \pm 15$ |
| $60\mu M$ TFV | $0.027 \pm 0.014$ | $4.2 \pm 0.73$ |
| $16nM$ ATV | $0.88 \pm 0.16$ | $3.2 \pm 0.088$ |

We then set out to investigate whether TFV and ATV could increase the probability of clearance of low inoculum infection, corresponding to *in vivo* exposure. We used 6.3*x*10^3^ viral copies (Materials and Methods), predicted to result in approximately 3 initial infected cells based on a regression of the number of infected cells versus input viral load (Figure 4A). Infection was initiated with the same cell-free viral dose for all conditions, and infected cells were cultured for 8 days in the presence of drugs. Steady state cell viability was maintained by passaging the culture using a 1:2 dilution of cells into fresh medium with drug. After 8 days, drug was washed away and infection amplified by growth for 6 days in the absence of drug, with passaging every 2 days, to detect the presence of any infected cells after drug treatment. For the no drug infection condition, it was suZcient to culture infection for 6 days to obtain a robust signal in terms of infected cells. The background level of GFP signal was estimated from uninfected cells (Materials and Methods). After amplification in the absence of drug for the TFV and ATV conditions, infection was either clearly visible or absent (Figure 4B).

We did not experimentally observe clearance of infection in the absence of drug. In the presence of TFV, clearance rose dramatically, with approximately three quarters of infections extinguished. In contrast, only a minor increase of infection clearance was observed with ATV (Figure 4C). Clearance with TFV was significantly higher relative to no drug and ATV, while ATV was not significantly different from no drug (*p* < 10^−5^ and *p* = 0.37, respectively, using bootstrap). Calculation of 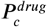 based on Eq. (5) using the measured values for *λ* and *R*_0_ for each drug condition showed a similar trend to the experimental values, with a higher clearance probability for each condition relative to the experimental results. Hence, Eq. (5) was able to predict the relative effectiveness of each drug to terminate infection in an *in vitro* model of PrEP.

**Figure 4.**
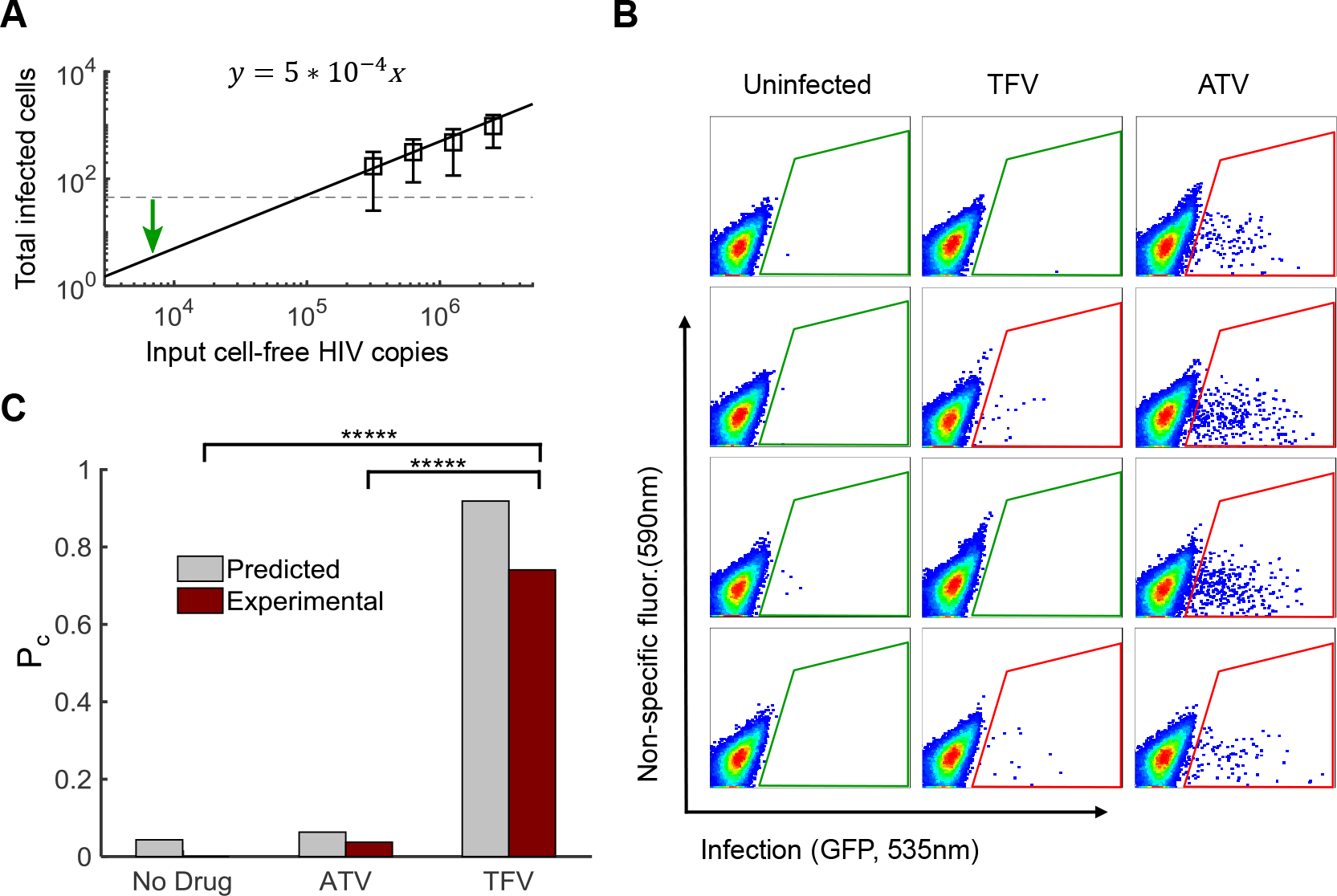
Probability of infection clearance depends on drug mechanism. (A) Determination of *λ*. The number of infected cells was measured using flow cytometry as a function of cell-free HIV RNA copies for four virus stock dilutions yielding a number of infected cells above the limit of detection after one infection cycle. Data was then fit using linear regression to determine the input viral dose for 3 infected cells. Mean ± std of 5 independent experiments. Dotted line is limit of detection. Green arrow marks number of HIV RNA copies used in the experiments. (B) Representative flow cytometry plots after 8 days of infection with the input cell-free virus in the presence of of TFV or ATV and further 6 days amplification of the same infection in the absence of drug. Each plot represents one independently cultured replicate of the experiment. Uninfected samples are shown in the left column, and infection in the presence of TFV or ATV is shown in the middle and right column respectively. X-axis is GFP fluorescence, y-axis is autofluorescence. The fraction of infected cells corresponds to the cells within the area outlined in green or red, with green indicating background GFP signal level as determined from the uninfected samples, and red indicating above background signal. (C) 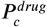 as predicted by Eq. (5) (gray bars) based on the measured drug effects on *R*_0_ and *λ*, and as experimentally measured (red bars). Presence of infection was assayed in 26 (no drug) or 27 (TFV and ATV) cell-free virus infected cell cultures from 4 independent experiments. Observed 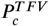 was significantly higher than 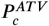 and 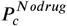 (*p* < 10^−5^ by bootstrap). 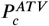 and 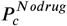 were not significantly different (*p* = 0.37 by bootstrap).

## Discussion

In this study we modeled and experimentally measured the clearance probability of initial HIV infection in the face of different inhibitor mechanisms: attenuation of infection frequency (TFV) or viral burst size from the infected cell (ATV). We chose drug concentrations where replication was reduced but not fully suppressed to investigate the effect of suboptimal HIV inhibition. Given that PrEP regimens seem to be more likely to be subject to variable adherence relative to treatment of established infection (*Van Damme et al.* (*2012*); *Marrazzo et al.* (*2015*); *Baeten et al.* (*2012*); *Choopanya et al.* (*2013*); *McCormack et al.* (*2016*); *Molina et al.* (*2015*); *Grant et al.* (*2014*)), sub-optimal inhibition is not only a function of the drug regimen but may be a feature of the approach This may make achieving *R*_0_ ⪅ 1 more difficult than with therapy for established infection. Under this set of conditions and in agreement with model predictions, only TFV, the inhibitor which was able to act before the first cell was infected, was able to increase the probability of HIV clearance. The prediction was based on two parameters which were readily measurable: *λ* is measured in one infection cycle with cell-free virus. *R*_0_ is measured using multiple viral cycles, but consists of relatively simple passaging of infection to ensure that uninfected target cells are not limiting. Prediction does require the calculation of the the Lambert function. This can be done by a wide range of algebra capable programs (in this study we used Matlab).

We show that the current PrEP approach utilizing drugs which target infection frequency is in fact the correct choice. However, the quantitative model presented here has utility beyond giving a rationale for what is already done. For example, ARVs with longer half-lives such as the integrase inhibitor cabotegravir may become future treatment modalities both as ongoing therapy and for PrEP (*Andrews and Heneine* (*2015*)). However, longer half-lives also result in the drug levels reaching steady state slower, as the time to reach 50 percent of the steady state concentration is one half-life (*Wagner* (*1975*)). Therefore, in early exposures post-administration, level of inhibition or drugs with long half-lives may be lower for initial infection relative to drugs of the same class but with shorter half-lives. This may mean that the calculated *P*_*c*_ may be different even among drugs which inhibit infection frequency and have similar effects on *R*_0_. In contrast, how fast a drug is able to act post-administration may have little effect in suppressing established infection. These considerations may also shape the effectiveness of future therapeutic approaches such as vaccines which require antigen based immune recognition: if *R*_0_ > 1 with the intervention, clearance would be unlikely if the immune mechanism is not rapid enough to prevent the first set of cells from being infected. Hence, under conditions where exposure may take place relatively rapidly post-PrEP dosing, the model presented here will be able to predict the relative PrEP utility of drugs of the same class at the level of cellular infection.

Factors in addition to drug effectiveness in inhibiting cellular infection may determine which drug or drug combination is used in PrEP. Among others, important considerations include drug side effects and interactions with other drugs (*Grant et al.* (*2010*); *Anderson et al.* (*2010*); *Choopanya et al.* (*2013*); *Koechlin et al.* (*2017*)), penetration to the initial site of infection (*Anderson et al.* (*2010*); *Patterson et al.* (*2011*); *Spinner et al.* (*2016*); *Murnane et al.* (*2013*); *Baeten et al.* (*2013*)), convenient dosing (*Grant et al.* (*2010*); *Anderson et al.* (*2010*); *Baeten et al.* (*2013*)), cost effectiveness (*Nichols et al.* (*2016*); *Anderson and Cooper* (*2009*); *Paltiel et al.* (*2009*); *Desai et al.* (*2008*); *Pretorius et al.* (*2010*); *Hallett et al.* (*2011*); *Gomez et al.* (*2012*)), and consequences that the use of the drug in PrEP will have on the transmission of resistant viral strains (*Supervie et al.* (*2010*, 2011); *Dolling et al.* (*2012*); *Dimitrov et al.* (*2013*); *Abbas et al.* (*2013*); *Saenz and Bonhoeffer* (*2013*); *Van De Vijver et al.* (*2013*); *Okano and Blower* (*2013*)). These additional factors need to be taken into account when designing a PrEP regimen.

The model output using the measured values for *λ* and *R*_0_ resulted in predicted probabilities of infection clearance which were higher than the experimentally observed clearance frequencies for all conditions. We speculate that the reason is an underestimation of the input number of infected cells *λ*. We measured *λ* one viral cycle after cell-free infection. If GFP expression in an infected cell was below threshold of detection at that time, the cell would be missed in the assay yet still amplify infection. Despite this, the relative effectiveness of each drug mechanism was clearly predicted by the model. Factors *in vivo* which may result in deviation from model predictions include transmission by cell-to-cell spread (*Sigal et al.* (*2011*); *Boullé et al.* (*2016*); *Jackson et al.* (*2018*); *Jolly et al.* (*2004*); *Duncan et al.* (*2014*); *Agosto et al.* (*2015*, 2018); *Law et al.* (*2016*); *Abela et al.* (*2012*); *Rudnicka et al.* (*2009*); *Sherer et al.* (*2007*); *Zhong et al.* (*2013*); *Del Portillo et al.* (*2011*); *Sowinski et al.* (*2008*); *Len et al.* (*2017*); *Baxter et al.* (*2014*); *Kim et al.* (*2018*)). This should reduce the effectiveness of PrEP since the drugs would only act on *R*_0_ and not on *λ*. A quantitative understanding such as presented here should detect deviations from the predictions made for cell-free infection, and thus allow a better understanding of any potential limits to PrEP.

HIV has been shown to rapidly seed a latent reservoir of infected cells (*Chun et al.* (*1997*a,b); *Finzi et al.* (*1999*); *Siliciano et al.* (*2003*); *Eriksson et al.* (*2013*)) which may also explain why it is necessary to block infection before the first cell is infected. However, at suboptimal inhibition, latent infection is not thought to play a role and infection is maintained by ongoing cycles of replication (*Sigal and Baltimore* (*2012*)). Evidence for this is the emergence of drug resistance with monotherapy (*Deeks et al.* (*1997*); *Molla et al.* (*1996*); *Wei et al.* (*2002*)). The current study provides a different rationale for why PrEP works best if the virus is inhibited before the first cellular infection, and enables a generalized evaluation of drugs to screen for the best candidates for this intervention.

## Materials and methods

### Inhibitors, viruses and cells

The following reagents were obtained through the AIDS Research and Reference Reagent Program, National Institute of Allergy and Infectious Diseases, National Institutes of Health: the antiretroviral drugs ATV and TFV; RevCEM cells from Y. Wu and J. Marsh; HIV-1 NL4-3 CCR5 tropic infectious molecular clone (pNL(AD8)) from E. Freed; pBABE.CCR5, from N. Landau. Cell-free virus was produced by transfection of HEK293 cells with pNL(AD8) using TransIT-LT1 (Mirus) transfection reagent. Supernatant containing released virus was harvested two days post-transfection and filtered through a 0.45 micron filter (GVS). The number of HIV RNA genomes in viral stocks was determined using the RealTime HIV-1 viral load test (Abbott Diagnostics). The RevCEM HIV infection GFP indicator cell line was modified as follows for experiments with the CCR5 tropic virus: The E7 clone was generated from RevCEM cells as described in *Boullé et al.* (*2016*). Briefly, the RevCEM cell line was sub-cloned by limiting dilution. Clones derived from single cells were expanded into duplicate 96-well plates, one optical and one standard tissue culture for continued growth. The optical plate was infected with NL4-3 and optical wells were scanned by microscopy to select clones with highest infection percentage by GFP expression. The clone E7 was selected based on >70 percent GFP positive cells upon infection, expanded from the uninfected replicate plate and frozen. To generate the CCR5 expressing B8 reporter clone, RevCEM-E7 cells were infected with the pBABE.CCR5 retroviral vector which stably expressed CCR5 under the LTR promoter. Cells were sub-cloned by limiting dilution. Clones derived from single cells were expanded into duplicate 96-well plates, one optical and one standard tissue culture for continued growth. The optical plate was infected with pNL(AD8) and wells were scanned by microscopy to select clones which maintained similar GFP expression to the parental REVCEME7 clonal cell line. The clone RevCEM-B8 was selected based on >70 percent GFP positive cells upon infection, expanded from the uninfected replicate plate, and frozen. Cell culture medium was complete RPMI 1640 supplemented with L-Glutamine, sodium pyruvate, HEPES, non-essential amino acids (Lonza), and 10 percent heat-inactivated FBS (Hyclone).

### Infection and flow cytometry

For determination of drug effect on *R*_0_ and *λ*, cells were infected with 2.5*x*10^7^ viral RNA copies in 2ml of cell culture containing 5*x*10^5^ cells/ml. Every 2 days, the uninfected and drug treated cell cultures were passaged at a split ratio of 1:2, where half the cell culture was removed and fresh media with drug (for TFV and ATV) or without drug (for uninfected) was added. For the no drug infection condition, 1:100 of the culture (20 *μ*l) was added to 2ml of fresh, uninfected cells at 5*x*10^5^ cells/ml. The removed fraction of cells was used to detect infection as follows: The number of infected cells was acquired every 2 days using 3 measurements of 30 second acquisitions per sample (approximately 50,000 cells/sample) with a FACSCaliber machine (BD Biosciences) using the 488nm laser line. Flow rate on the machine was measured at each time-point, and acquisition time was multiplied by flow rate to obtain the number of infected cells per ml. For experiments measuring 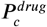, 200 *μ*l of cells at a density of 5*x*10^5^ cells/ml were infected with 6.3*x*10^3^ viral RNA copies. For cell cultures with either no infection or containing ATV or TFV, passaging conditions were the same as for the experiments used to determine drug effect on *R*_0_ except that no culture was removed. Instead, new media with drug was added for the TFV and ATV conditions, and new media with no drug was added for the no infection condition. The infection volume therefore doubled every 2 days, and the cell culture was transferred to larger volume wells to preserve a constant surface to volume ratio. After 8 days (4 passages), cells were spun down, washed once in medium with no drug, and resuspended at 5*x*10^5^ cells/ml in fresh medium with no drug. Cells were then further passaged in the absence of drug for 6 days (3 passages) using a 1:2 dilution every 2 days to amplify any infection in the culture. For infection in the absence of drug, cells were passaged for 6 days (3 passages) using a 1:2 dilution every 2 days with fresh medium without removing any of the cell culture. The number of infected cells was acquired at the end of the experiment (14 days post-infection for the uninfected, TFV and ATV conditions and 6 days for the no drug infection condition) as 30 second acquisitions per sample with a FACSCaliber machine as above. Results were analyzed using FlowJo 10.0.8 software. The background frequency of positive cells was determined by acquiring uninfected samples (n=17 from 4 independent experiments). A sample was scored as infected if the number of GFP positive cells was greater than that in the highest background samples (4 positive cells per 30 second acquisition).

### Statistical significance by bootstrap

The vector of TFV results where infection was scored as either present (1) or absent (0) at each position in the vector, was resampled with replacement using 10^5^ iterations to determine the number of iterations which resulted in a vector of 26 1’s (no drug) or 26 of 27 1’s (ATV).

### Derivation of the mathematical model

The probability that one cell with a burst of *κ* virions will infect *m* cells and not infect *κ* – *m* cells is:

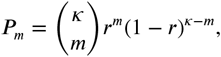

where *r* id the infection probability per cell and 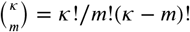. We decline the generating function for this probability distribution:

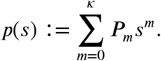

Using the binomial formula 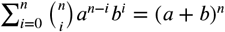, we obtain

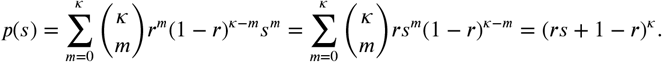

For *R* > 1, the extinction probability of infection is given by the smallest root of *p*(*s*) = *s* (*Grimmett and Stirzaker* (*2001*)), that is:

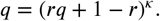

In the biologically relevant case where *κ* is large and *r* is small, with *R*_0_ finite, we approximate the solution (*rs* + 1 – *r*)^*κ*^ = *e*^*κ* ln(*rs*+1–*r*)^ = *e*^*κ* ln(1+*r*(*s*–1))^ ≃ *e*^*κr*(*s*–1)^. Therefore,

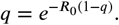

The solution for *q* to this equation is presented as Eq. (1).

## Acknowledgments

AS thanks Israel Michael Sigal for extensive discussions, and Deenan Pillay, Mark Siedner, and Brenda Tipper for comments on the manuscript.

